# The effect of a free radical scavenger on oxidation stress in partial bladder outlet obstruction and its relief in a rat model

**DOI:** 10.1101/348151

**Authors:** Min Soo Choo, SongZhe Piao, Seung-June Oh

**Affiliations:** Department of Urology, Hallym University Dongtan Sacred Heart Hospital, Hwaseong, Korea; Department of Urology, Seoul National University Hospital, Seoul, Korea; Department of Urology, Post-Doctoral Research Center, Yanbian University Hospital, Yanji, China

**Author notes:** Corresponding author: Seung-June Oh, MD, PhD Department of Urology, Seoul National University Hospital, 101 Daehak-ro, Jongno-gu, Seoul 03080, Korea / /.

**Keywords:** urinary bladder neck obstruction, prostate hyperplasia, urinary bladder, overactive, oxidative stress, free radical scavengers, receptors, adrenergic, beta-3

## Abstract

**AIMS:** To investigate the effect of a free radical scavenger (tempol) after relief of partial bladder outlet obstruction (pBOO) on bladder function in a rat model.

**METHODS:** pBOO was induced in 50 eight-week-old female Sprague-Dawley rats and relieved 3 weeks later. The rats were divided randomly into 5 groups: sham-operated, tempol-treated for 1 week (Treat-1w) or 3 weeks (Treat-3w), and no treatment for 1 week (nonTreat-1w) or 3 weeks (nonTreat-3w). Awaken cystometrograms were obtained 1 or 3 weeks after relief according to the grouping. The bladders were isolated and weighed. H&E, Masson’s trichrome and TUNEL staining were used to analyze histological changes. The oxidative stress assessed using malondialdehyde. The expression of beta-3 adrenoreceptor was examined by Western blotting.

**RESULTS:** The tempol-treated groups exhibited a significant decrease in the number of IDCs per voiding cycle (nonTreat-1w vs. Treat-1w, 1.18±0.82 vs. 0.36±0.40, P=0.010; nonTreat-3w vs. Treat-3w, 1.51±0.69 vs. 0.23±0.25, P=0.002). The thickness and collagen fiber deposition of the detrusor muscle layer was significantly decreased in the treated groups. Apoptosis detected was mainly observed in the urothelial cell layer, although the rate of apoptosis was significantly decreased in the treated groups (48.9±3.36% vs. 32.7±11.10%, P=0.024; 25.8±4.67% vs. 15.7±9.83%, P=0.314). The tempol-treated groups showed significant decreases in the MDA concentrations at both 1 and 3 weeks after relief. The expression of the beta-3 adrenoreceptor was increased in the tempol-treated rats.

**CONCLUSIONS:** Ischemic reperfusion injury after relief of pBOO caused histological and functional changes in the bladder. Free radical scavenger treatment prevented this oxidative stress.

## Introduction

Bladder outlet obstruction (BOO) occurs due to a variety of causes regardless of age, including the posterior urothelial valve in children, urethral stricture in adults, and benign prostatic hyperplasia (BPH) in the elderly [1]. Although the most common cause in the clinic is BPH in older men, BOO occurs in women with several anatomical and/or functional etiologies, including pelvic organ prolapse, Skene’s gland cyst, primary bladder neck obstruction, and detrusor external sphincter dyssynergia [2]. BOO is one of the most important clinical problems and can cause overactive bladder, urinary incontinence, urinary tract infection, vesicoureteral reflux, hydronephrosis, and renal insufficiency through chronic urinary retention [3].

Surgical treatment of BOO may temporarily exacerbate urgency related to detrusor overactivity (DO) and even cause transient urinary urge incontinence [4]. Newly developed urinary urge incontinence after a HoLEP procedure has been reported in 7.1-44.0% of patients [5]. This de novo urge incontinence causes significant stress and anxiety not only for the patients but also for the surgeons. In a recent prospective study of persistent storage symptoms after successful relief of BOO, urodynamic DO was persistent in approximately 40% of the patients [6].

Ischemia reperfusion injury (I/R injury) has been suggested to be a cause of post-operative bladder dysfunction in BPH patients [7]. In patients with chronic BOO due to BPH, a chronic ischemic status is induced in the inner bladder wall due to persistent high intravesical pressure [8]. In this condition, relief of chronic obstruction causes reperfusion and reoxygenation, resulting in the generation of reactive oxygen species (ROS), which cause more severe oxidative damage and cell apoptosis in the bladder wall [9].

Efforts to prevent I/R injuries have been made in various medical fields. Several antioxidants have been used to reduce I/R injury in patients with ischemic diseases, such as coronary artery disease or stroke, and in the transplantation field. For instance, L-alanyl-glutamine attenuates I/R injury in liver transplantation patients [10]. Some antioxidants, such as CoQ10, beta carotene, lycopene, quercetin, resveratrol, vitamin C and vitamin E, have shown preventive and therapeutic benefits for different forms of CVD [11]. However, no studies have investigated methods to prevent I/R injury after BPH surgery. The effects of I/R injury after BPH surgery have only recently drawn researchers’ attention.

In this study, we investigated the effects of oxidative stress caused by I/R injury after relief of partial BOO (pBOO) on bladder functions using a rat model and evaluated the preventive effect of free radical scavengers on bladder functions after relief of pBOO.

## Materials and Methods

### Animals and study design

Seven-week-old female Sprague-Dawley rats weighing between 220 and 250 g were used in this study. The rats were housed in a vivarium with free access to food and water in a light-controlled room with a diurnal cycle for 1 week prior to surgery. After surgery, the animals were caged individually and maintained under the same conditions. All animal handling and treatment procedures followed the Guide for the Care and Use of Laboratory Animals of the National Institutes of Health and were approved by the Institutional Animal Care and Use Committee of our institution, which is an AAALAC-accredited facility.

Fifty rats were randomly divided into five groups (n=10 rats per group). The first group consisted of control sham-operated rats (control group). The other four groups underwent urethral constriction to procedurally induce pBOO, followed by a reversal operation after 3 weeks. The second group underwent a cystometrogram (CMG) without treatment for one week after the reversal operation (unTreat-1w group). The third group underwent a CMG without treatment for 3 weeks after the reversal (unTreat-3w group). The fourth group underwent a CMG with tempol treatment for 1 week after the reversal (Treat-1w group). The fifth group underwent a CMG with tempol treatment for 3 weeks after the reversal (Treat-3w group). The rats in the treated group received tempol (Sigma-Aldrich, St. Louis, MO, USA) by gavage at a dose of 1.5 mmol/kg/day dissolved in water three times per day [12].

### Induction of pBOO

Anesthesia was induced by ketamine/xylazine (15 mg/kg and 5 mg/kg; intramuscular injection) and maintained with isoflurane (1-3%). After shaving the skin, asepsis was attained with a 10% povidone-iodine solution while the rat was in a dorsal recumbent position. The bladder was approached through a lower midline abdominal incision. After exposing the proximal urethra, a steel rod 0.9-mm in diameter was placed around the urethra. The bladder neck was ligated using a 3-0 silk, and the steel rod was removed. The bladder was repositioned, and the abdominal wall was closed. In the sham operation group, the bladder neck was very loosely ligated to avoid inducing any obstruction. Three weeks after inducing pBOO, each rat underwent a reversal operation that removed the ligation in the same manner. We measured the body weight of each rat before each procedure.

### Procedures for intravesical and intra-abdominal catheter implantation

The catheter implantation procedures were performed 2 days before the functional evaluation. Polyethylene catheters (PE-50; Clay-Adams, Parsippany, NJ, USA) with a cuff were inserted into the dome of the bladder through a lower abdominal incision with a purse-string suture and simultaneously placed on the posterior side of the bladder. The balloon and catheter were filled with distilled water, and the distal end of the catheter was sealed. Both catheters were tunneled subcutaneously to the skin of the back and anchored with a silk ligature. The free end of the catheter was sealed. Each rat was housed individually after the procedures and maintained in the manner described above.

### Functional evaluation

CMGs were performed on awakened, unanesthetized, and unrestrained rats in metabolic cages after a minimum of 2 days of recovery from catheterization. The bladder catheter was connected via a 3-way stopcock to a pressure transducer (Research Grade Blood Pressure Transducer; Harvard Apparatus, Holliston, MA, USA) and a microinjection pump (PHD22/2000 pump; Harvard Apparatus). Another pressure transducer was connected to an intra-abdominal balloon catheter to independently record the intra-abdominal pressure (IAP). The micturition volumes were recorded with a fluid collector connected to a force displacement transducer (Research Grade Isometric Transducer; Harvard Apparatus). Room-temperature saline was infused into the bladders at a rate of 0.4 mL/min. Pressures and micturition volumes were recorded continuously with a computerized system (PowerLab, ADInstruments, Colorado Springs, CO, USA) at a sampling rate of 50 Hz. Non-voiding contractions (NVCs) during the filling phase were defined as an increment of intravesical pressure that exceeded 2 cmH_2_O from the baseline without simultaneous changes in IAP and without fluid expulsion from the bladder [13].

When the bladder contractions became stable, at least five micturition cycles were recorded for each rat. The following CMG parameters were measured, and the mean value of each variable was calculated for the analysis: basal pressure (the lowest pressure during filling), threshold pressure (the pressure immediately before the initiation of micturition), peak micturition pressure (the maximum pressure during micturition), micturition interval (the interval between micturition contractions), micturition volume, micturition duration (the time from initiation to finish of one micturition cycle), post-voided residual volume (the remaining urine after voiding), bladder capacity (the infused volume immediately before the initiation of micturition), bladder compliance (calculated by dividing the micturition volume by the difference between resting and threshold pressures), and the frequency of NVCs (per micturition cycle).

### Histology and immunohistochemistry (IHC)

The rats were euthanized after completion of the functional study. The whole bladder was extirpated and weighed. A portion of the bladder was fixed in a 4% formaldehyde solution for the histological evaluation, and some portions were stored at −80°C for the IHC evaluation.

After obtaining 4-μm serial sections of the paraffin-embedded material, the bladder tissue sections were stained with hematoxylin and eosin or Masson’s trichrome stain. The thickness of the detrusor muscle layer was evaluated in 10 randomly selected hematoxylin and eosin-stained sections. Collagen deposition was measured in 10 high-power (×400) fields from randomly selected Masson’s trichrome-stained sections. Photomicrographs were obtained using a digitalized miscroscopic image system (Nikon Eclipse 80i microscope and Nikon Digital Slight DS-U3; Nikon, Tokyo, Japan). The images were analyzed using the Adobe and ImageJ software (http://rsb.info.nih.gov/ij/).

To detect apoptosis, the terminal deoxynucleotidyl transferase-mediated dUTP-biotin nick end labeling (TUNEL) method was used. In each slide, 10 high-power (×400) fields were randomly selected, and the apoptotic index was expressed as the percentage of apoptotic cells relative to the number of total cells in a given area (nonapoptotic nuclei plus apoptotic cells).

### Measurement of malondialdehyde (MDA) in the bladder

The MDA level in the bladder tissue, which is an index of oxidative stress, was determined using a commercially available kit according to the manufacturer’s instructions. The MDA concentrations were normalized using the protein content (NWLSSTM Malondialdehyde Assay, Northwest Life Science Specialties, LLC, Vancouver, WA, USA).

### Western blotting analysis of beta-receptor expression

The membranes were blocked with 5% skim milk for 1 hr at room temperature and incubated overnight at 4°C with primary antibodies against the β3 adrenergic receptors (1:1000, ab59685, Abcam), followed by incubation with appropriate horseradish peroxidase-linked secondary antibodies (1:4000) for 2 hr at room temperature. The bands on the blots were visualized using an enhanced chemiluminescence system (Amersham Bioscience, Buchinghamshire, UK), and densitometric analysis of the Western blots was conducted using VisionWorks LS, version 6.7.1.

### Statistical analysis

All data are expressed as the mean and standard error of the mean (SEM). The collected data were analyzed using Student’s t-test or the Mann-Whitney U test depending on whether the data followed a Gaussian distribution. A two-tailed P<0.05 was considered significant. The statistical analyses were performed using IBM SPSS for Windows, Version 24.0 (IBM Inc., Armonk, NY, USA).

## Results

### Body and bladder weights

No significant differences in body weight were found between the sham-operated and the pBOO-induced groups 1 week after relief of pBOO. However, the urinary bladder weight was significantly increased in the pBOO-induced groups compared with the weights in the sham group 1 week after the reversal operation (28.2±8.3 g vs 12.5±0.7 g, P<0.001). The ratio of bladder weight to total body weight was significantly higher in the pBOO-induced groups than in the sham group. No differences were found in body weight, bladder weight, and the ratio of bladder weight to body weight between the treated and untreated groups.

### Comparison of cystometric parameters

Significant differences were found in all cystometric parameters except the micturition duration, including the basal pressure, threshold pressure, peak micturition pressure, micturition interval, and voided volume, between the sham and the pBOO-induced groups. A significant difference was found in the NVC incidence between the tempol-treated and the untreated groups. The number of NVCs per voiding cycle was significantly decreased in the tempol-treated groups compared to the numbers in both the 1- and 3-week untreated groups (Treat-1w vs. unTreat-1w, 0.36±0.40 vs. 1.18±0.82, P=0.010; Treat-3w vs. unTreat-3w, 0.23±0.25 vs. 1.51±0.69, P=0.002). However, no other differences in the cystometric parameters were associated with tempol treatment (Table 1).

### Histological findings

Histologically, a thickened bladder wall was observed in the pBOO-induced bladder specimens. This finding was mainly due to hypertrophy of the detrusor muscle layer. In the untreated group, the thickness of the detrusor muscle layer decreased significantly after 3 weeks compared to 1 week of relief. However, in the tempol-treated group, no significant difference in the thickness of the detrusor muscle layer was observed between 1 and 3 weeks after relief.

In the comparison with the untreated group, a significant decrease was observed in the detrusor muscle thickness in the treated group after both 1 and 3 weeks of treatment (Treat-1w vs. unTreat-1w, 776.45±140.78 vs. 1164.17±190.58, P<0.001; Treat-3w vs. unTreat-3w, 726.26±162.76 vs. 905.82±161.16, P=0.043) (Figure 1).

**Figure 1.**
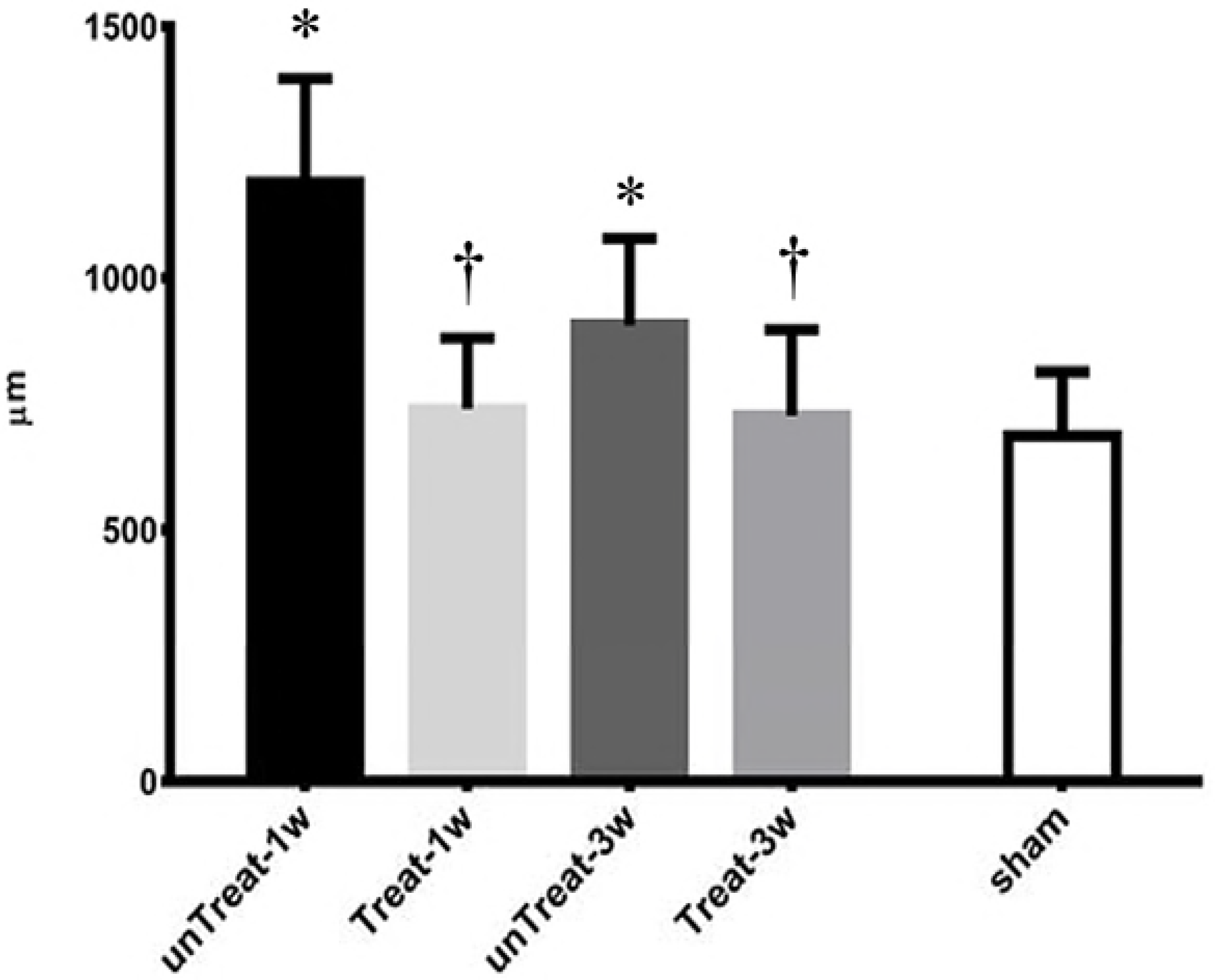
Changes in the detrusor muscle thickness after pBOO and its relief for 1 wk and 3 wks based on tempol treatment. Representative microscophs of a thin section of the bladder wall (magnification 100 ×). Bar graphs show the quantitative image analysis. Results are expressed as mean ± standard error of the mean. ⓐ untreated for 1 week; ⓑ tempol-treated for 1 week; ⓒ untreated for 3 weeks; ⓓ tempol-treated for 3 weeks; ⓔ sham; *P<0.05 versus sham; ^†^P<0.05 versus untreated group in the same period.

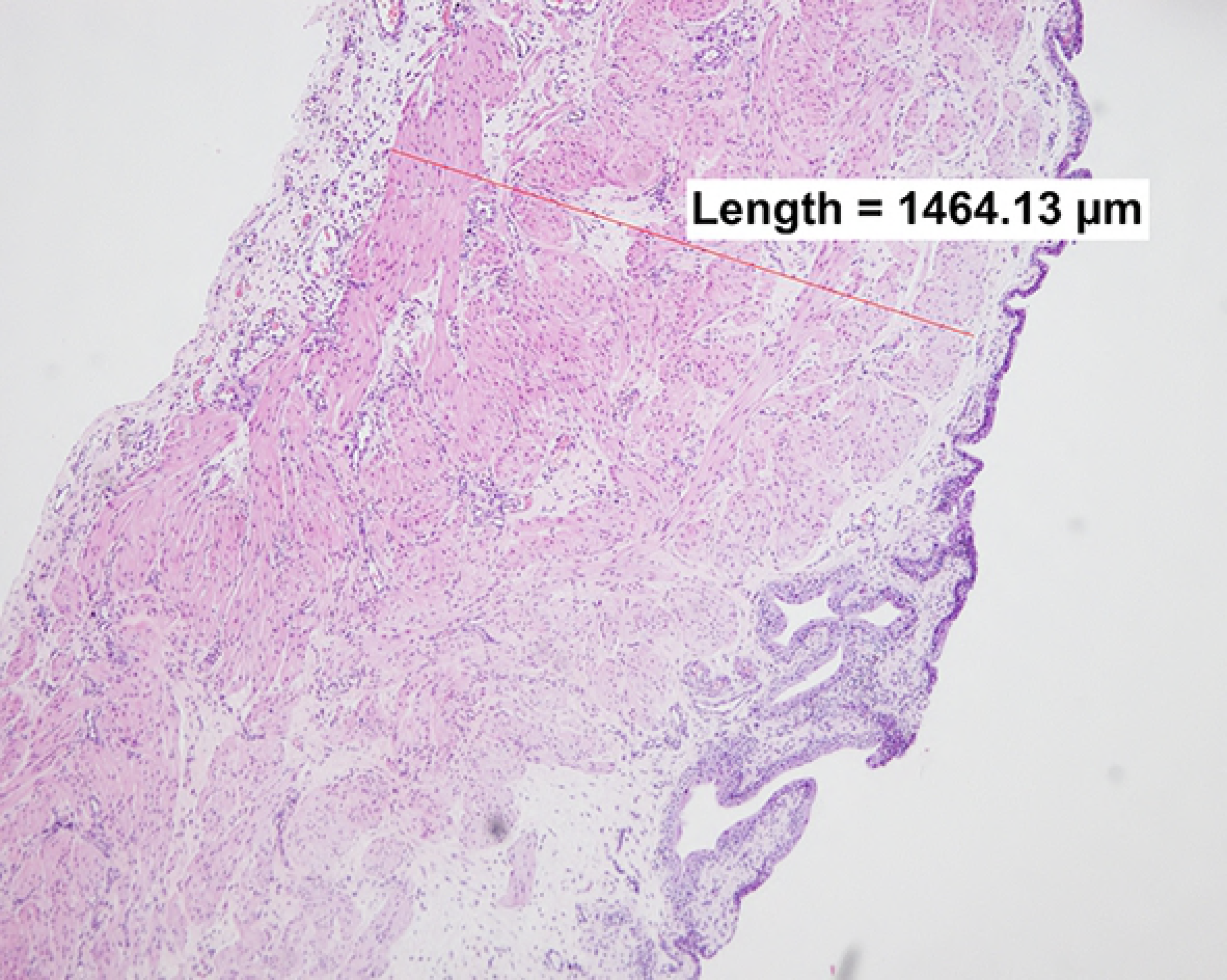

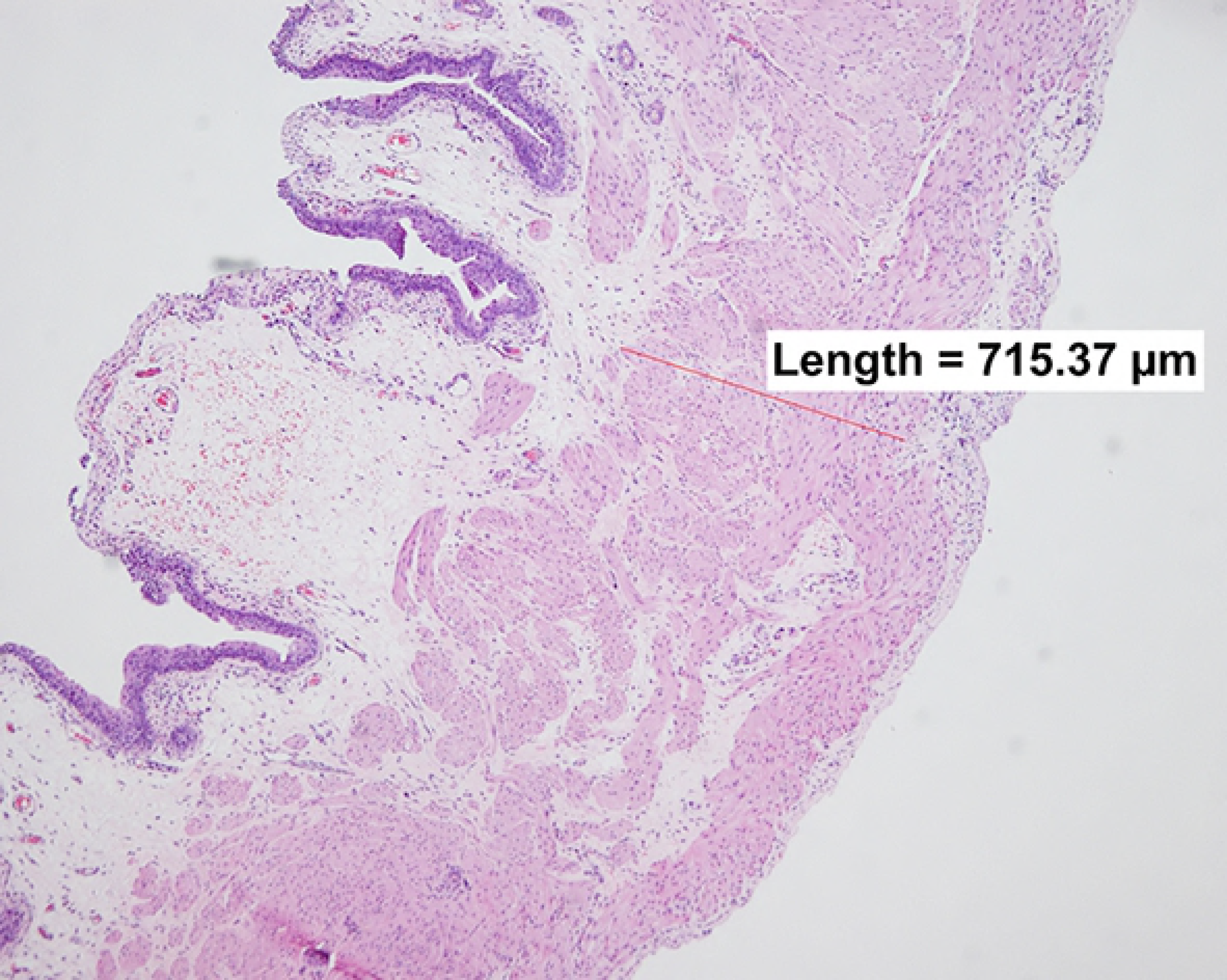

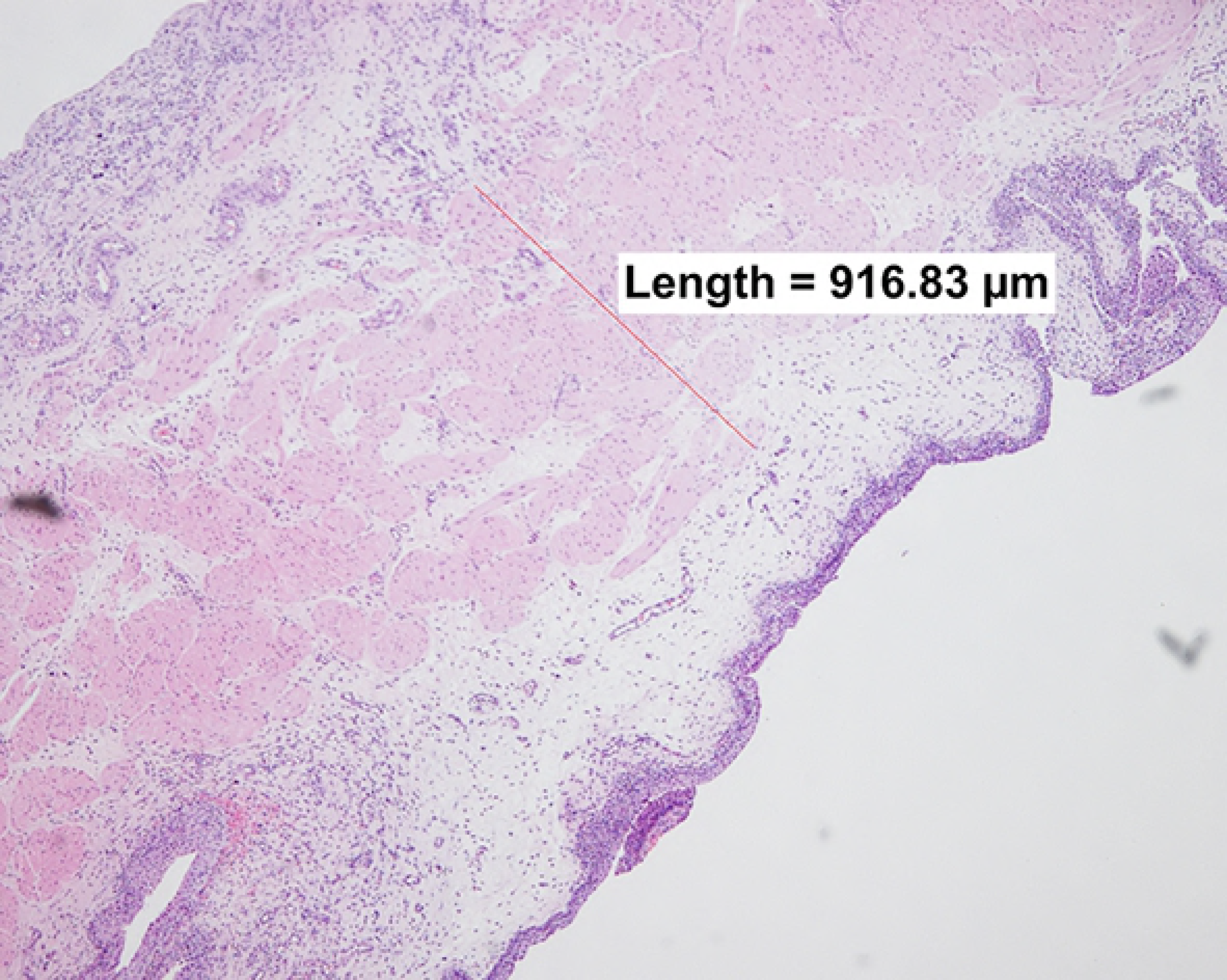

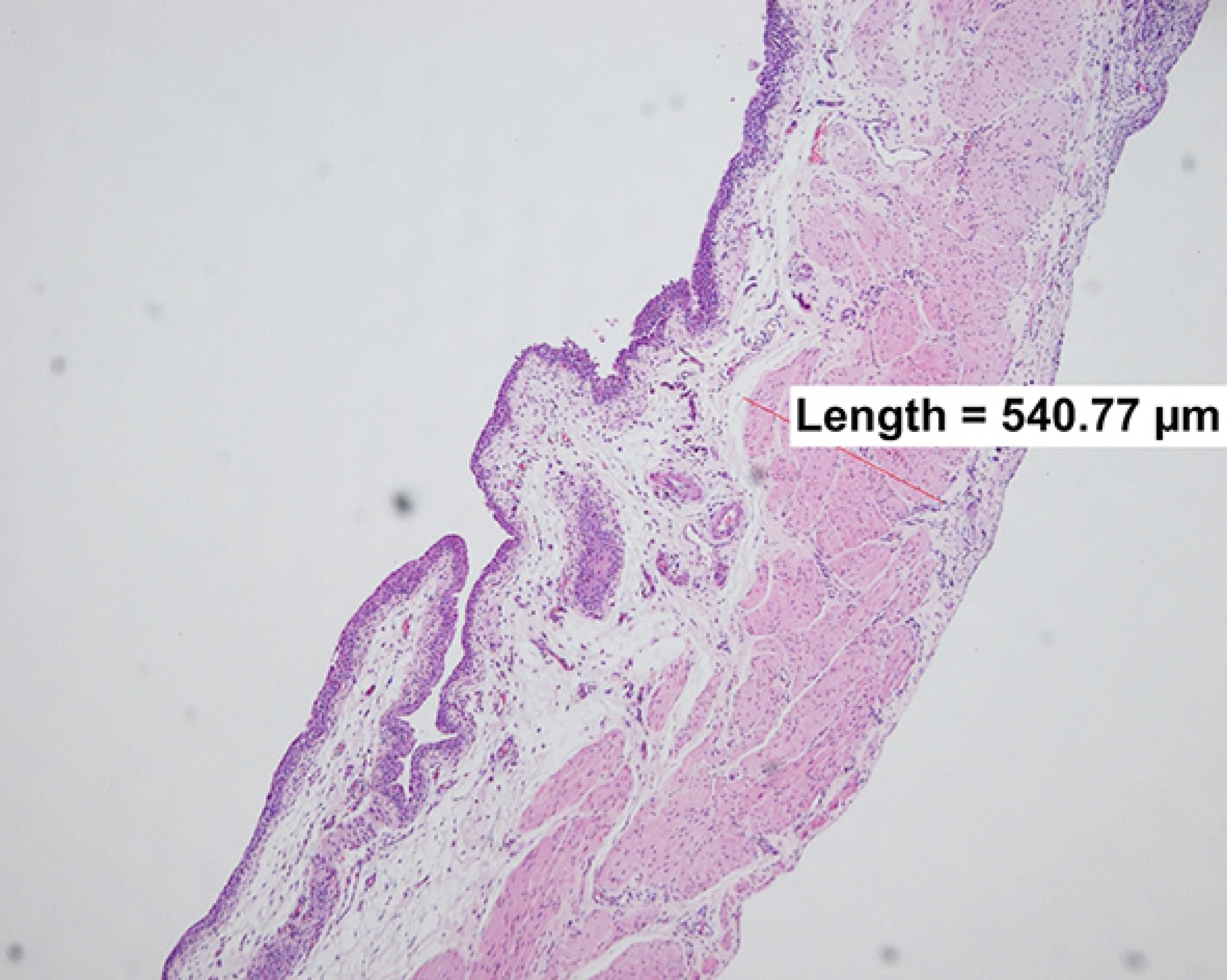

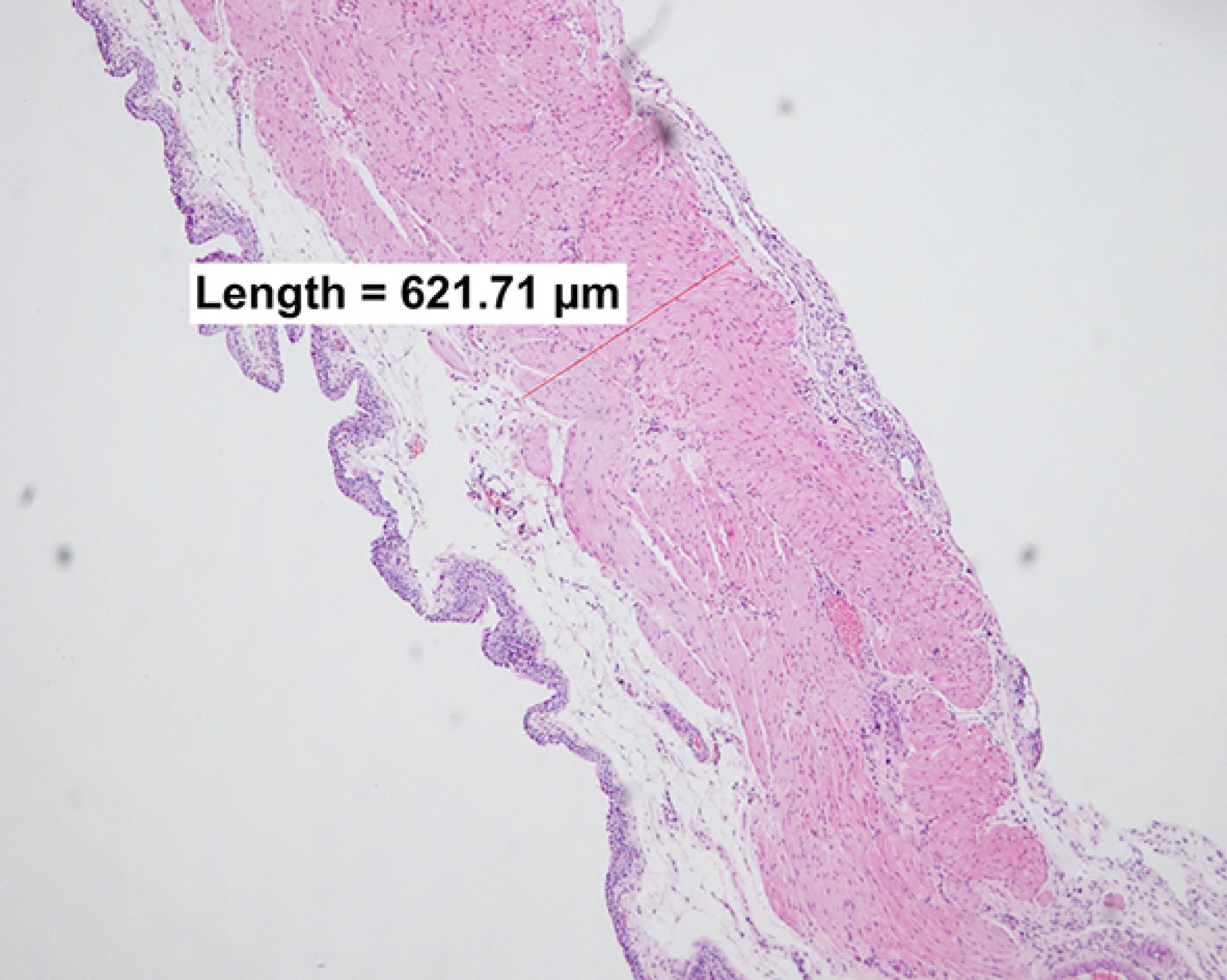

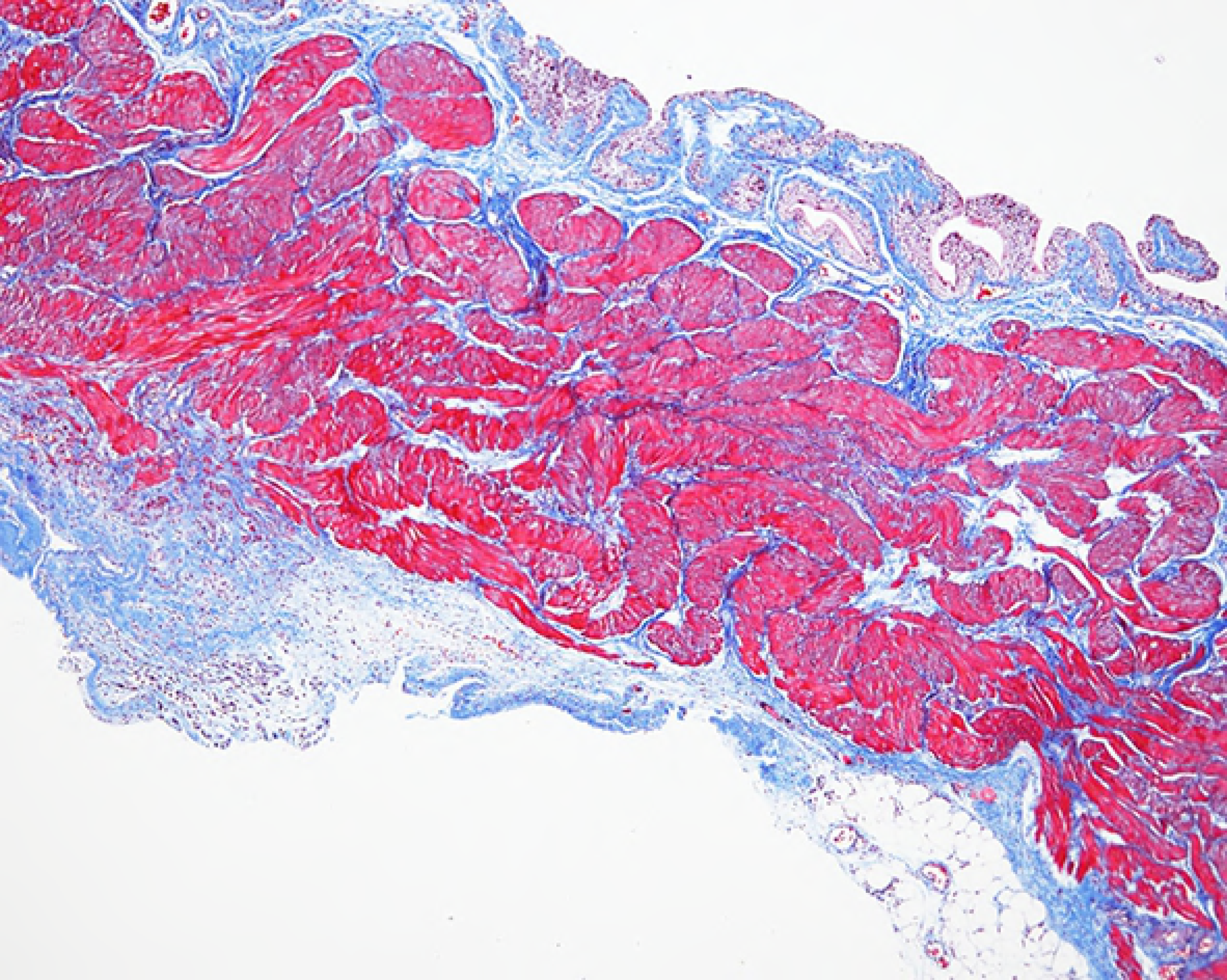

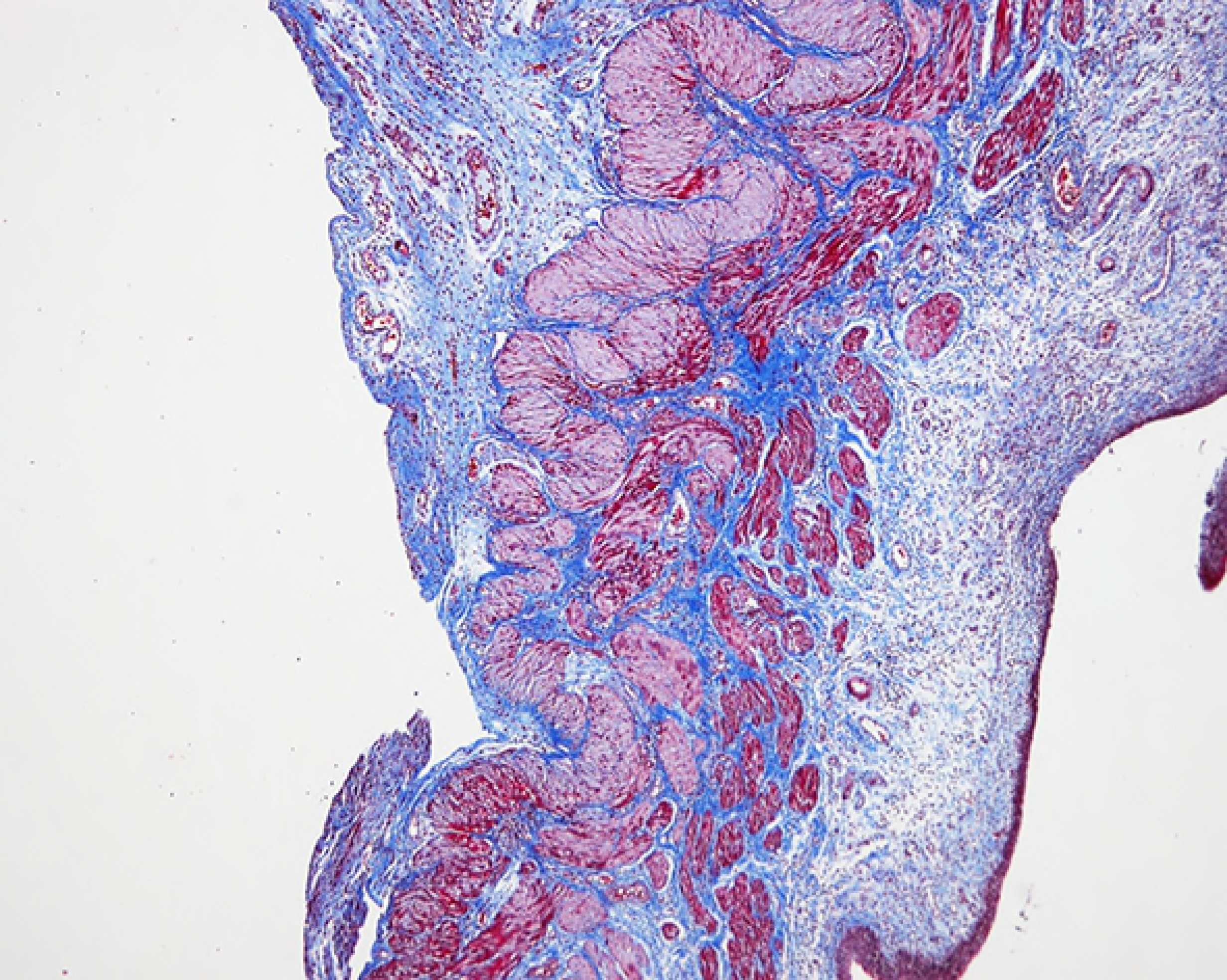

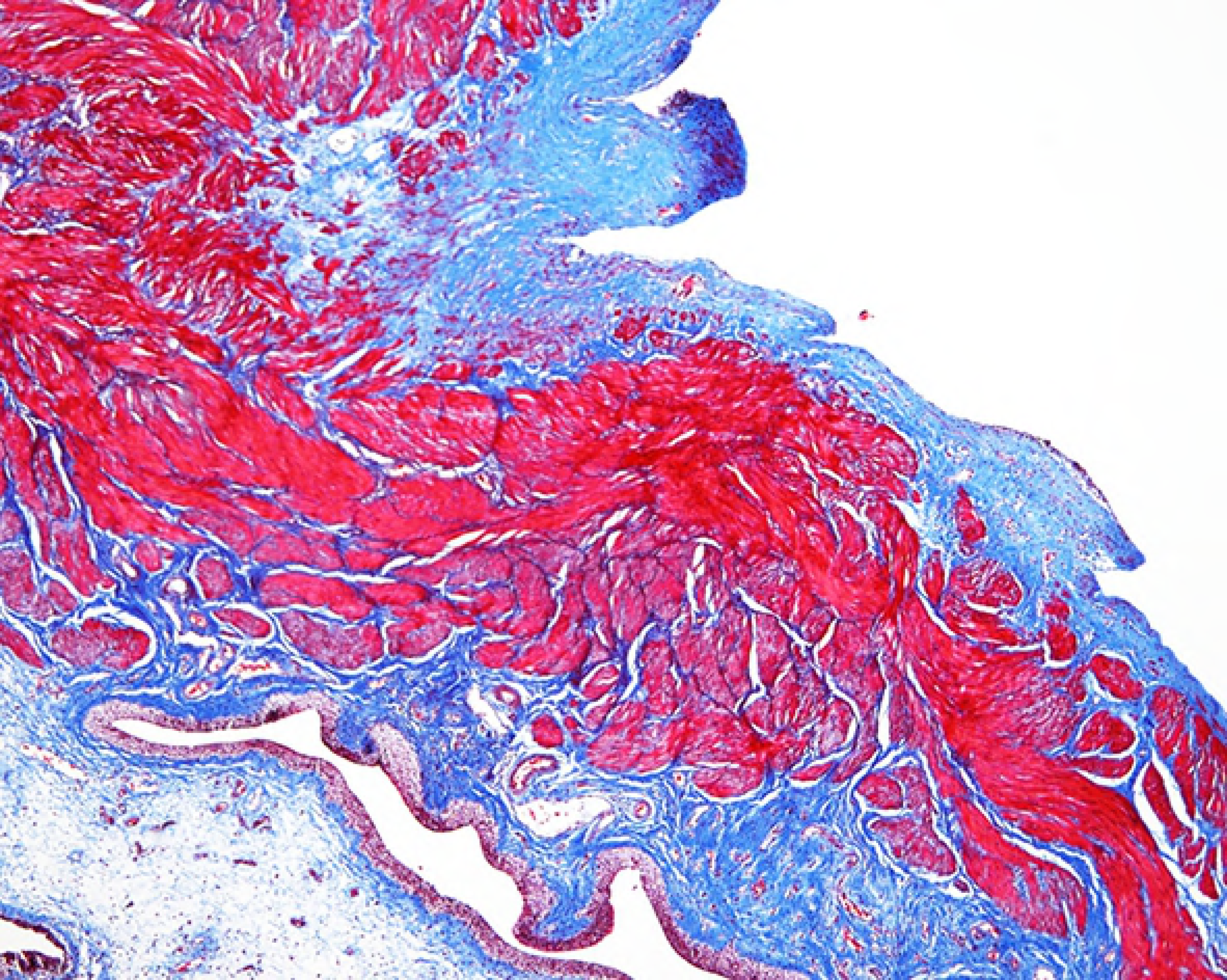

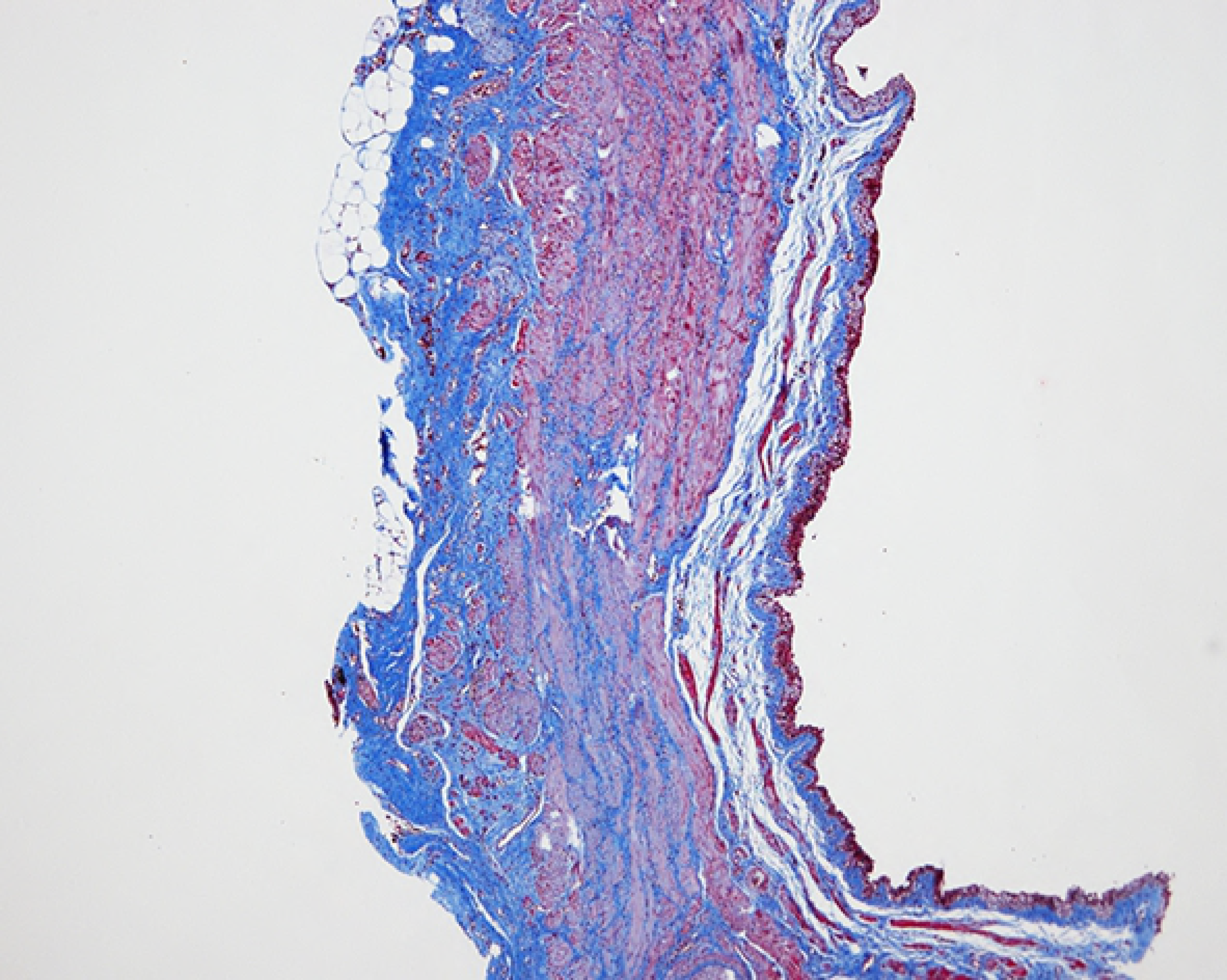

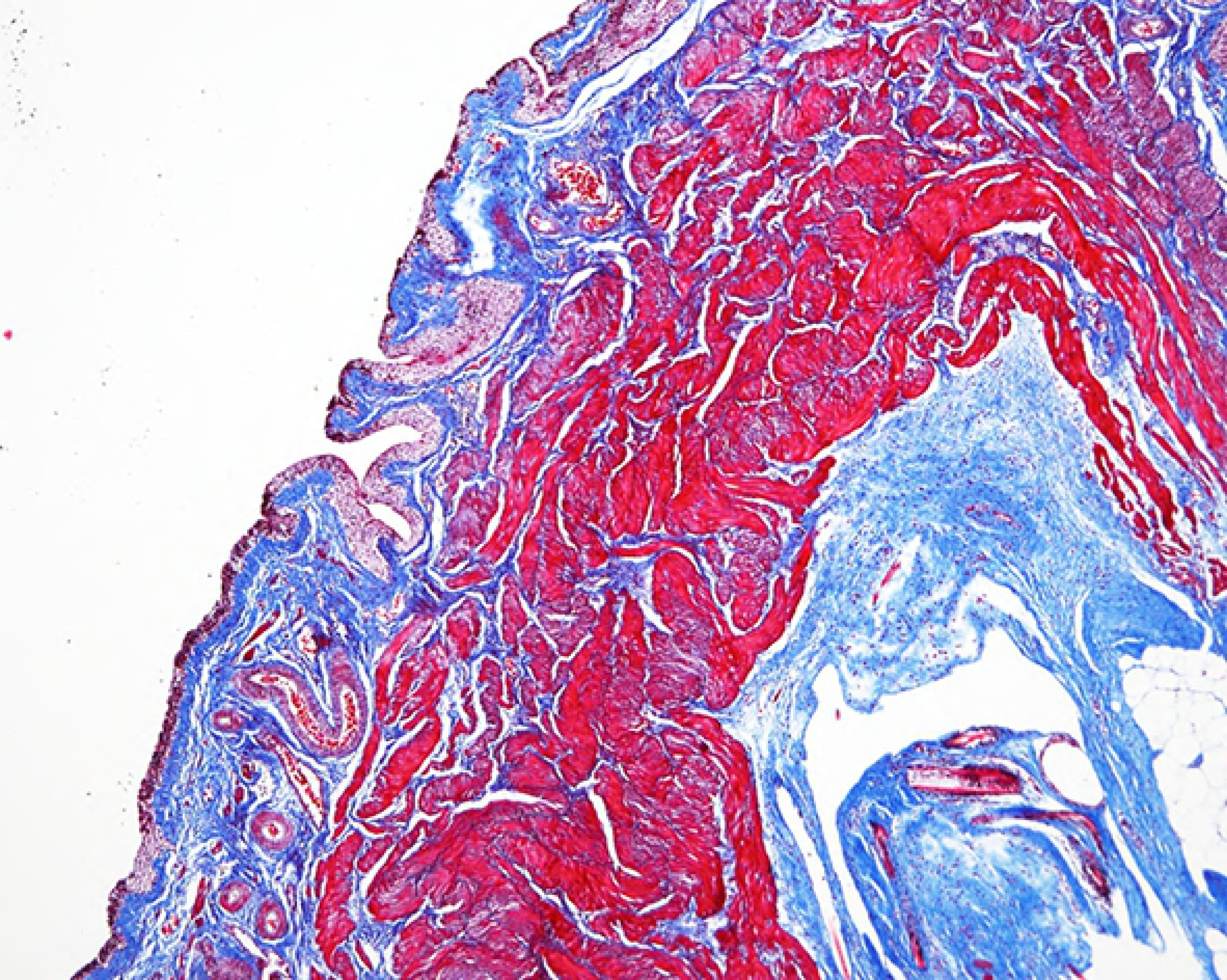

In the Masson’s trichrome-stained sections, tempol treatment reduced the deposition of collagen fibers in the lamina propria and detrusor muscle layer compared with the analysis in the untreated rats. The ratio of collagen to smooth muscle was significantly decreased in the treated rats at 1 week after relief, but not statistically at 3 weeks (Figure 2).

**Figure 2.**
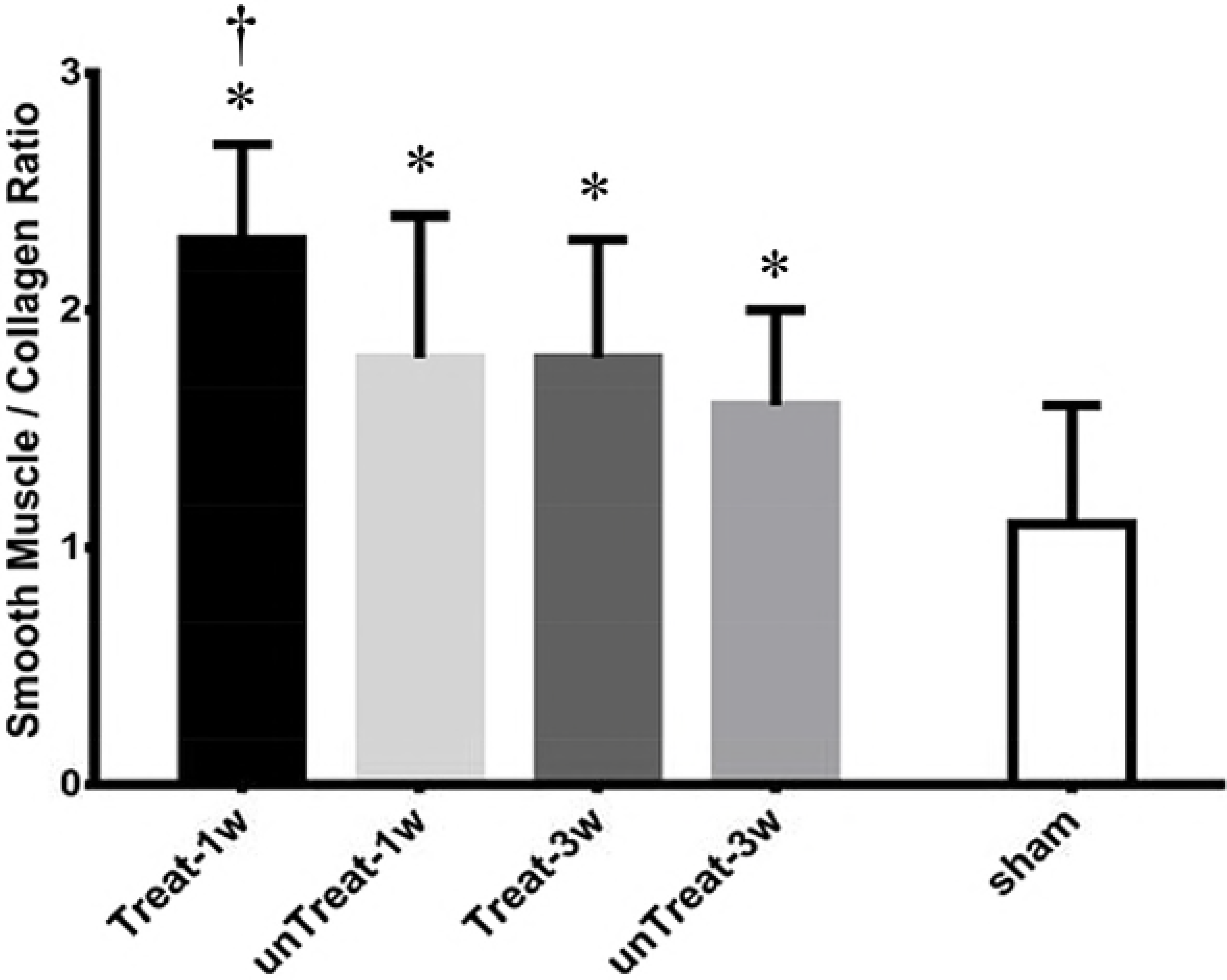
Masson’s trichrome staining: Representative microscophs show collagen with blue staining and the muscle with purple staining (magnification 20 ×). Bar graphs show the quantitative image analysis. Results are expressed as mean ± standard error of the mean. ⓐ untreated for 1 week; ⓑ tempol-treated for 1 week; ⓒ untreated for 3 weeks; ⓓ tempol-treated for 3 wks; ⓔ sham; *P<0.05 versus sham; ^†^P<0.05 versus untreated group in the same period.

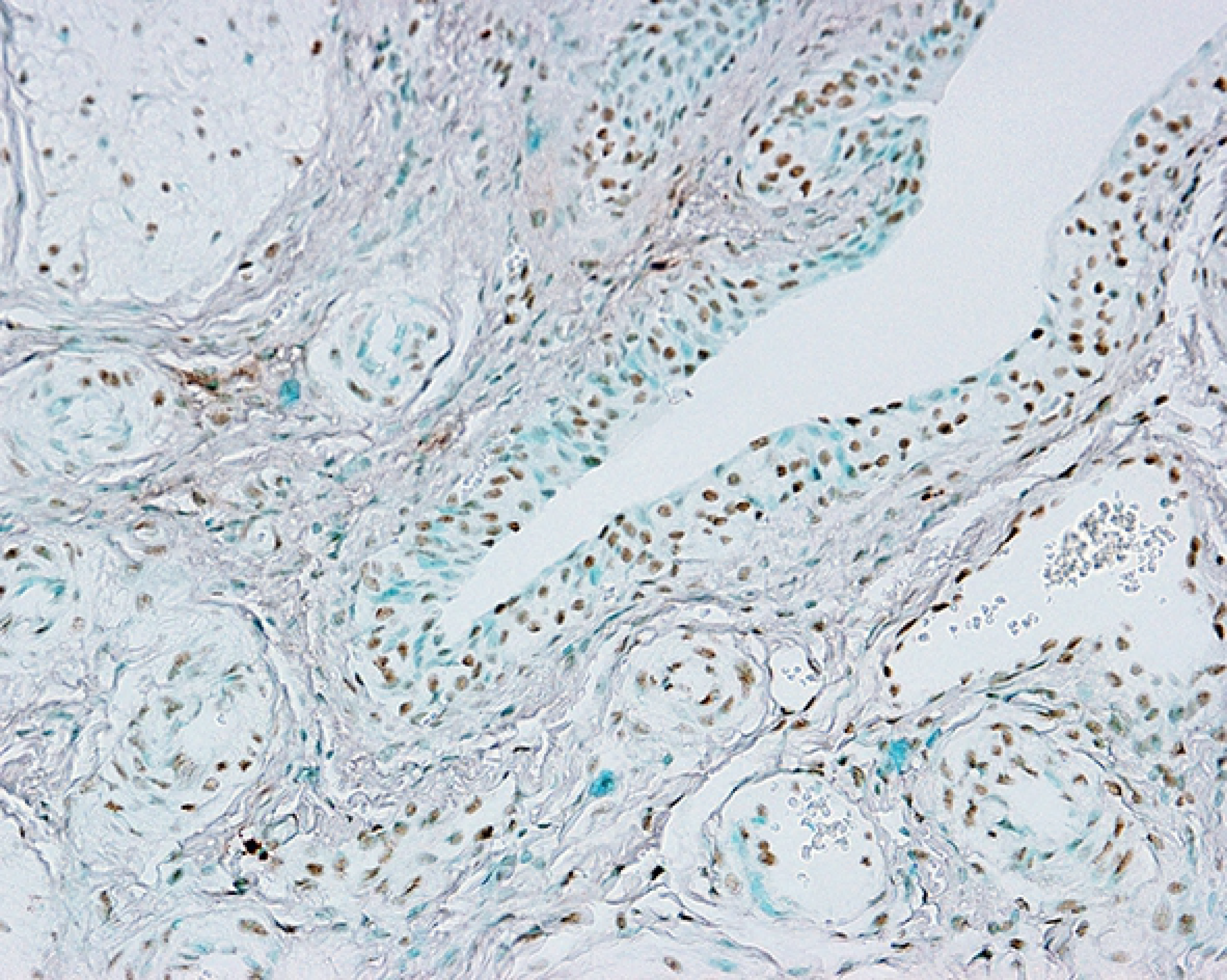

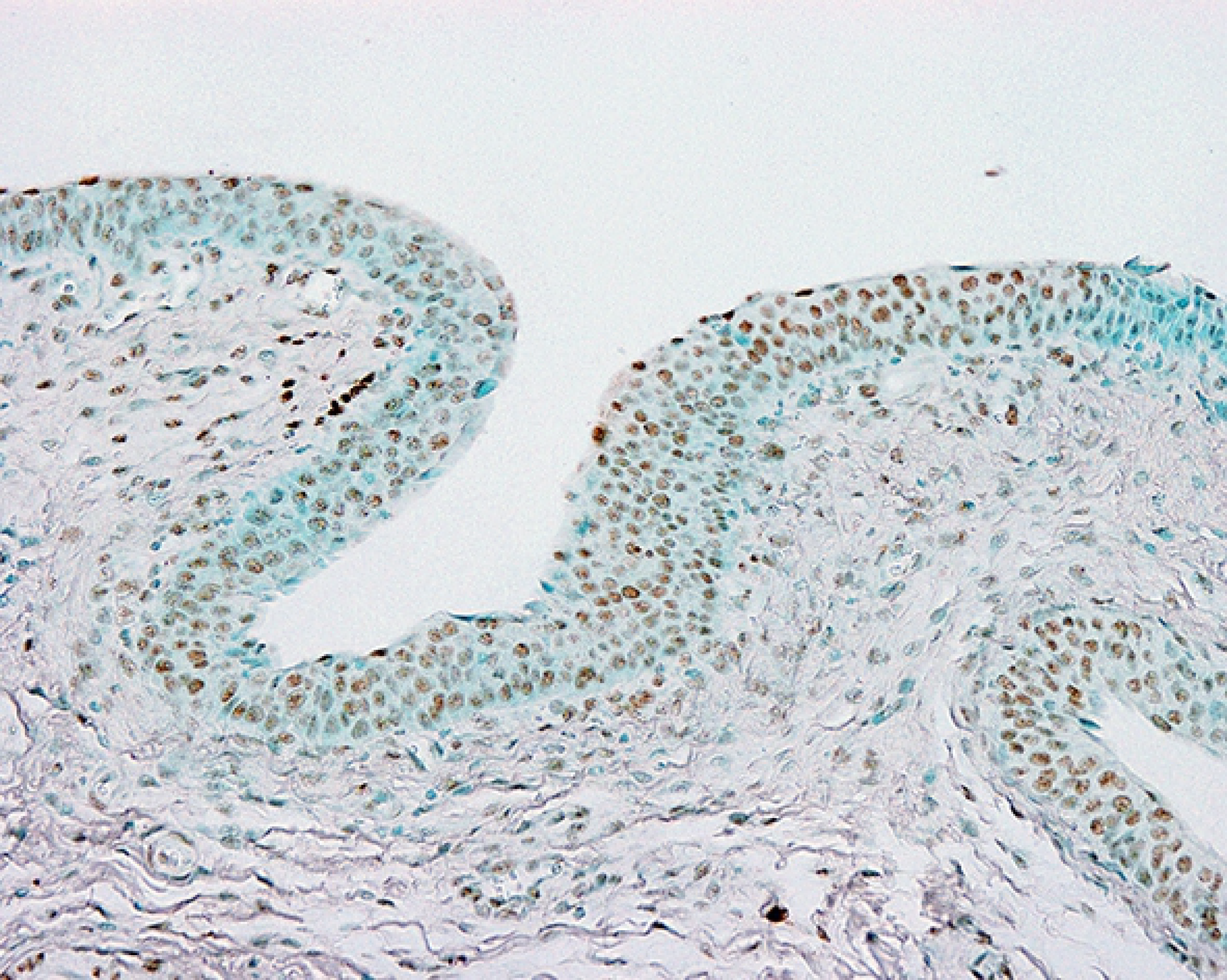

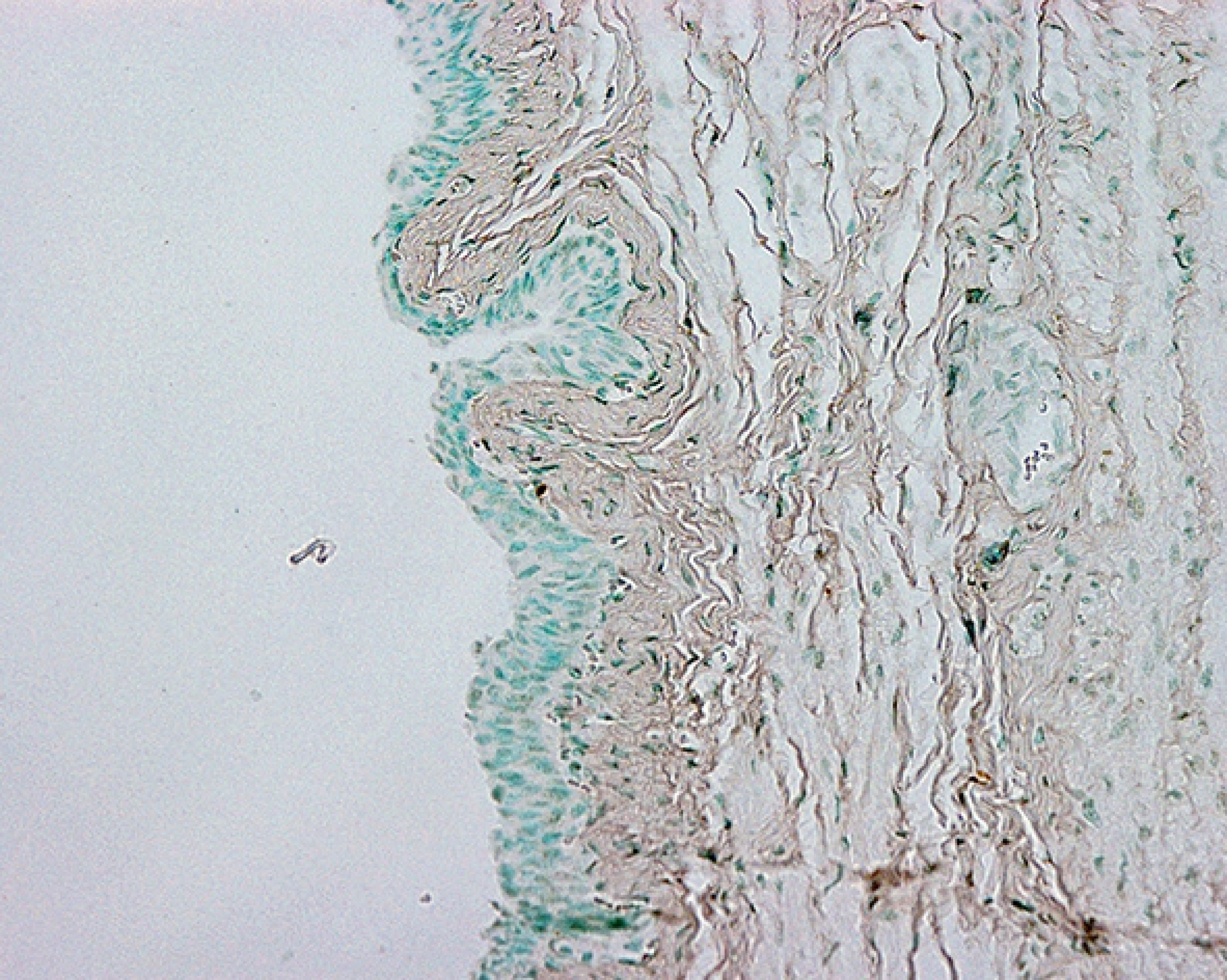

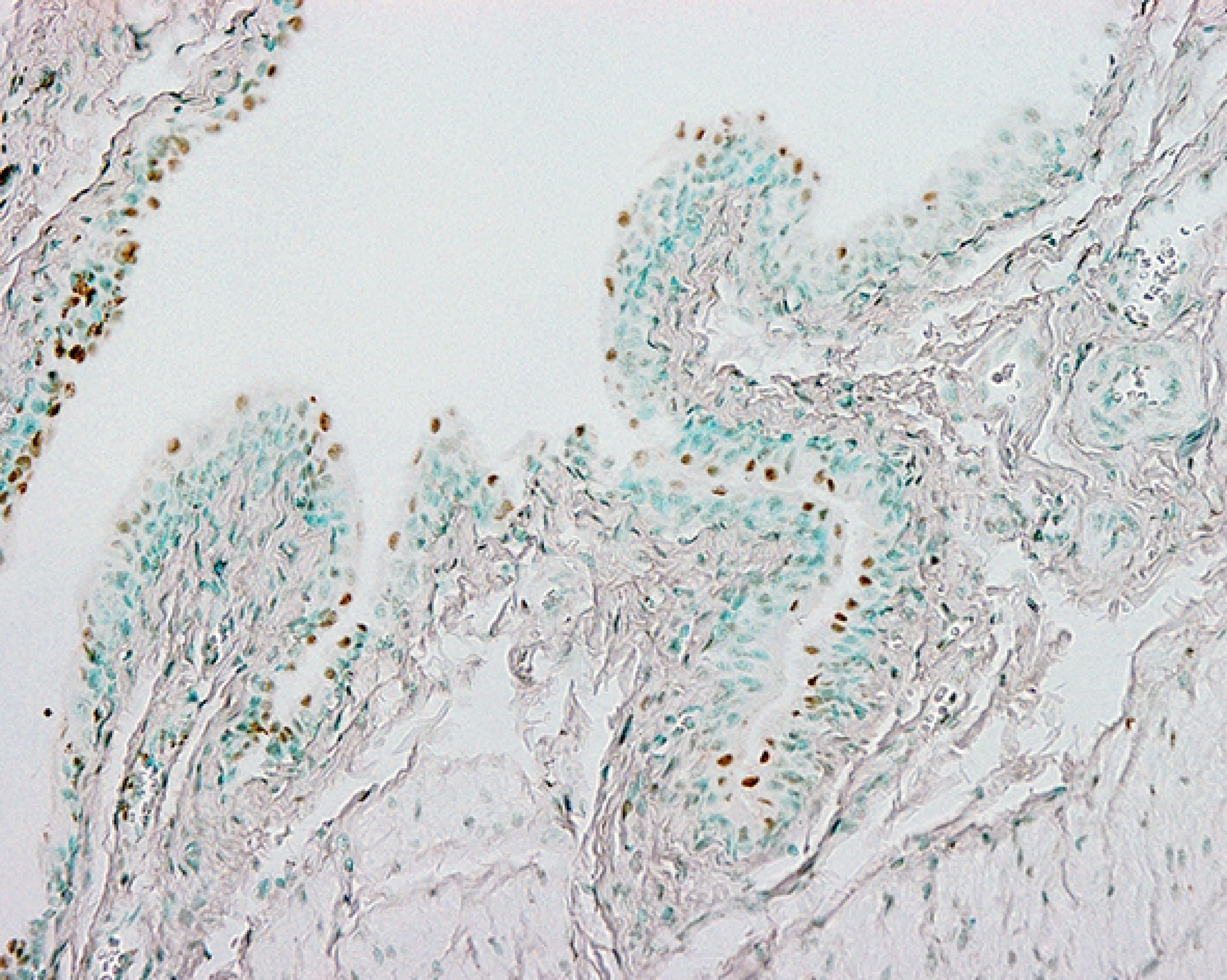

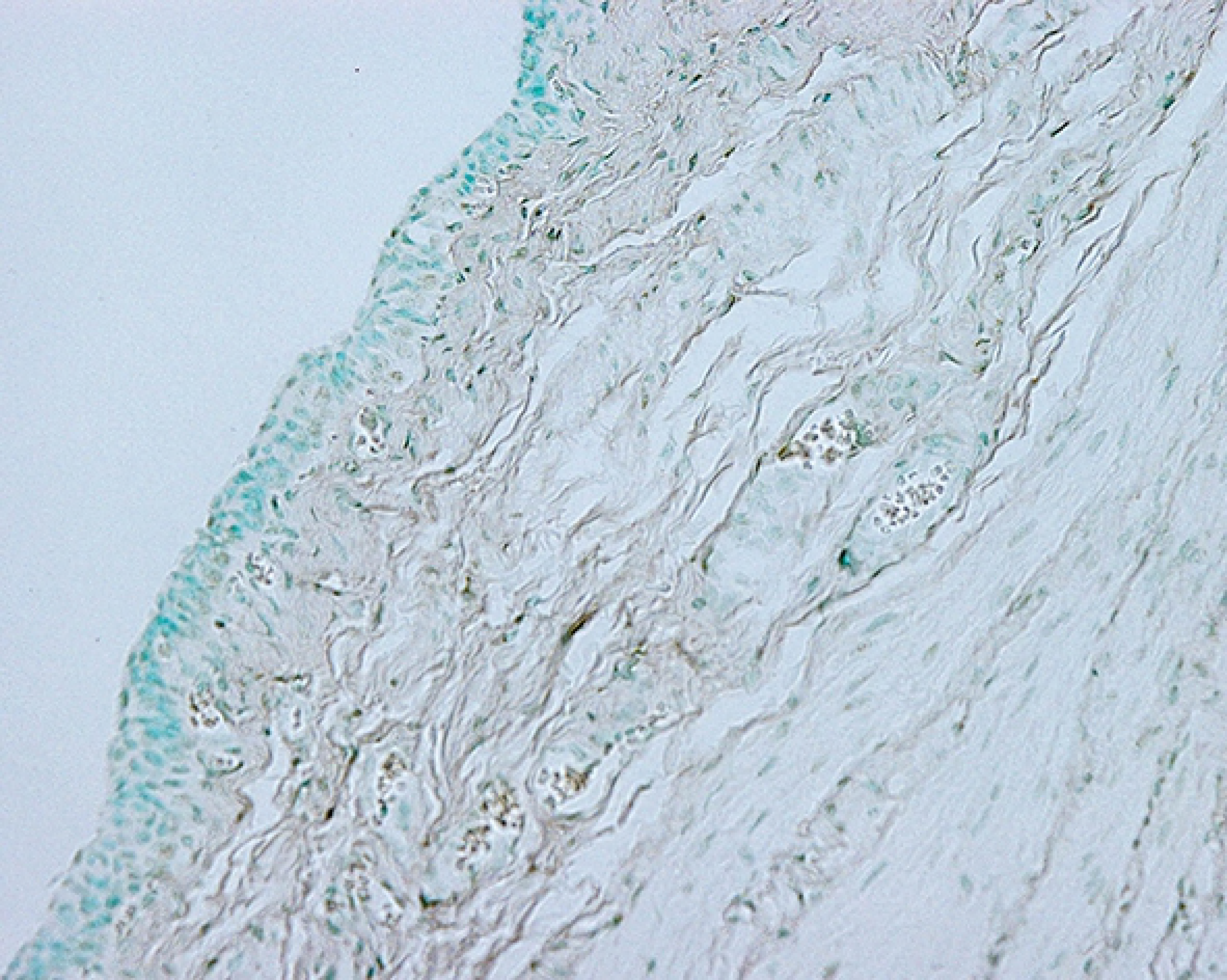

### TUNEL findings

TUNEL-positive cells were observed mainly in the urothelium. The numbers of TUNEL-positive cells were significantly decreased in the tempol-treated groups. This inhibitory effect was observed at both 1 and 3 weeks after relief (Treat-1w vs. unTreat-1w, 32.7±11.10% vs. 48.9±3.36%, P=0.024; Treat-3w vs. unTreat-3w, 15.7±9.83% vs. 25.8±4.67%, P=0.314) (Figure 3).

**Figure 3.**
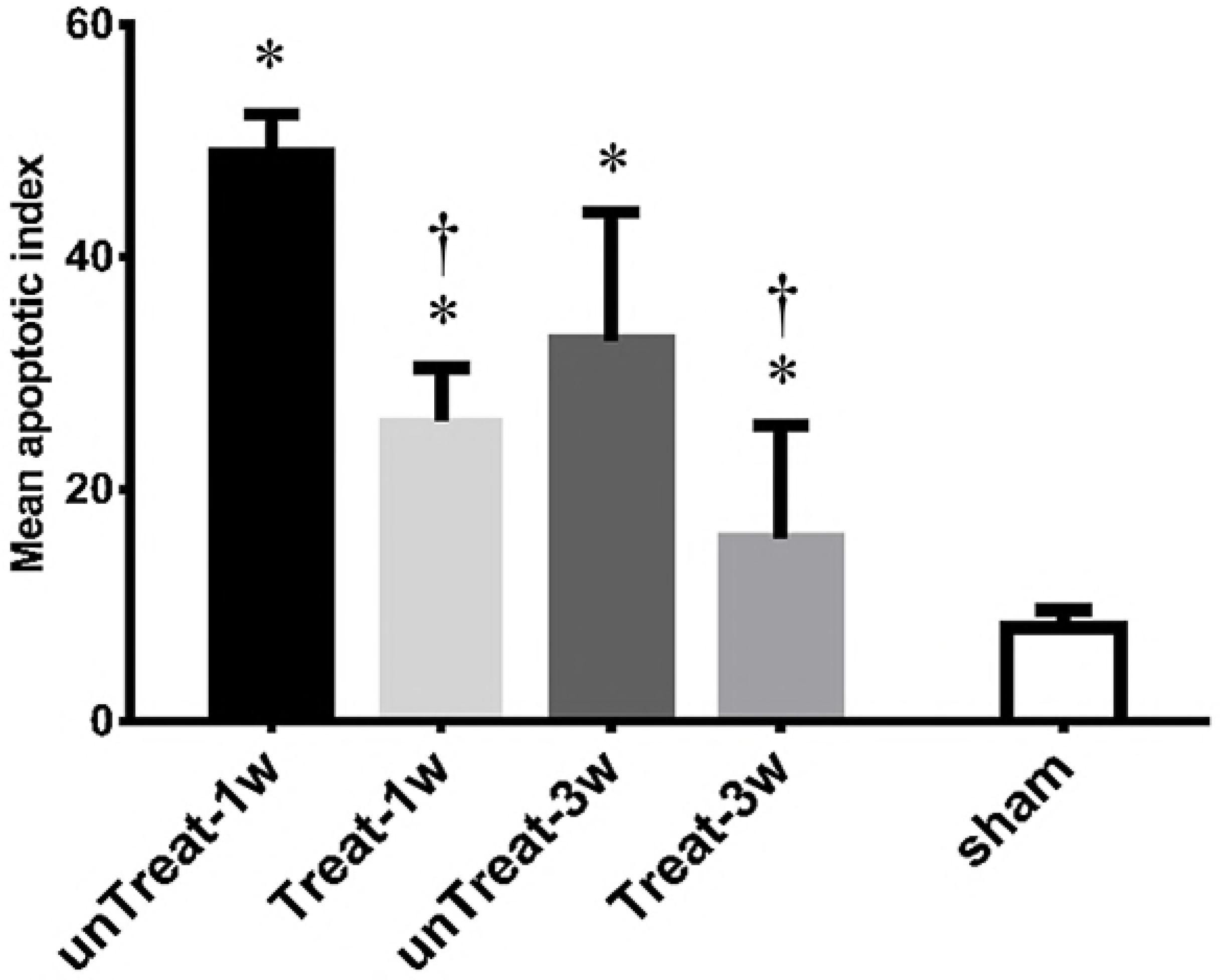
Detection of apoptosis: Representative microscophs show TUNEL-positive cells as black-brown cells mainly localized in the bladder urothelium (magnification 400 ×). Bar graphs show the quantitative image analysis. The apoptotic index represents the percentage of apoptotic cells within the total number of cells in a given area. Results are expressed as mean ± standard error of the mean. ⓐ untreated for 1 week; ⓑ tempol-treated for 1 week; ⓒ untreated for 3 weeks; ⓓ tempol-treated for 3 weeks; ⓔ sham; *P<0.05 versus sham; ^†^P<0.05 versus untreated group in the same period.

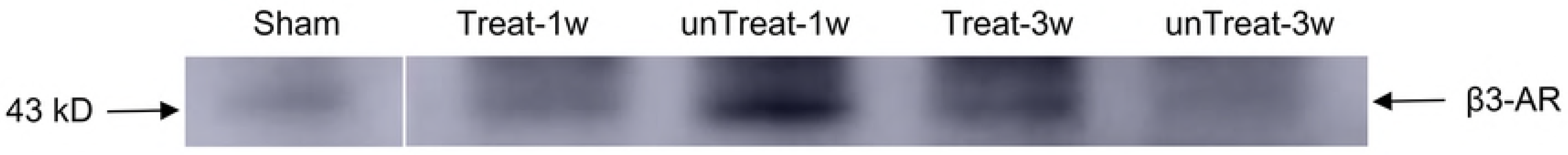

### MDA

The tempol-treated groups showed significant decreases in the MDA concentrations at both 1 and 3 weeks after relief compared to the concentrations in the untreated groups (Treat-1w vs. unTreat-1w, 0.66±0.07 vs. 0.76±0.08, P=0.021; Treat-3w vs. unTreat-3w, 0.48±0.06 vs. 0.58±0.10, P=0.030).

### Western blotting

The abundance of the beta-3 adrenoreceptor was increased in the tempol-treated rats 3 week after relief compared to the expression levels in the untreated rats. However, no significant difference in the abundance of the beta-3 adrenoceptor was observed 1 weeks after relief between the treated and untreated rats (Figure 4).

**Figure 4.**
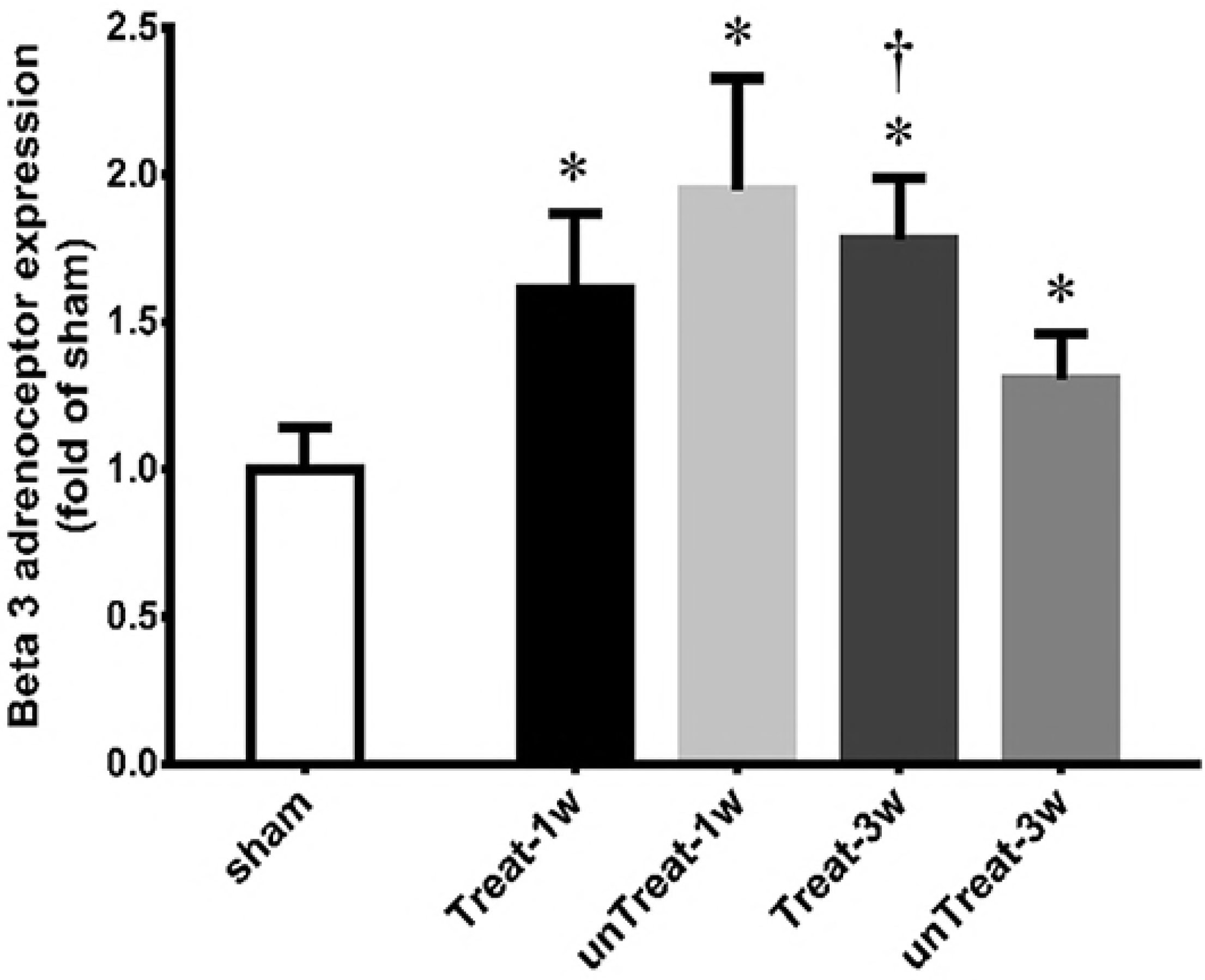
Western blot analysis: Representative Western blot shows expression of beta-3 receptor protein in the bladder, respectively. Bar graphs show the quantitative analysis for beta-3 receptor protein expression. Data are expressed as folds of corresponding expression in sham-operated rats. Results are expressed as mean ± standard error of the mean. *P<0.05 versus sham; ^†^P<0.05 versus untreated group in the same period.

## Discussion

In this study, we demonstrated in an animal model that antioxidants, such as tempol, prevented bladder I/R injury after relief of pBOO. The preventive effect lasted up to three weeks. The preventive effect of antioxidants on I/R injury was shown to reduce apoptosis mainly in the mucosal layer and to restore muscle hypertrophy more rapidly in the smooth muscle layer. This effect resulted in a decrease in NVCs in cystometry that was presumably related to beta-3 adrenoreceptor expression.

pBOO causes a chronic ischemic status in the bladder. High pressure due to sustained overdistention can induce a reduction of blood flow to the bladder wall [14]. Additionally, pBOO increases the thickness of the bladder wall through hypertrophy of the detrusor smooth muscle and deposition of collagen tissues, resulting in a reduction of microvascular blood perfusion [15]. In addition, rapid reperfusion due to relief of pBOO in chronically ischemic bladders leads to the generation of free radicals and rapid oxidative stress. Relief of pBOO resolves the intravesical high pressure, although resolving the bladder wall thickness takes at least 3 weeks, and the ongoing reduction of microvascular blood perfusion lasts for a considerable period. The increased metabolic demand on the hypertrophied detrusor can produce more severe damage in the bladder when combined with a reduction in microvascular blood perfusion [16]. The decompensated bladder also causes substantial residual urine, and overdistension of the bladder persists for a considerable period of time, resulting in sustained I/R injury after relief. In our study, this I/R injury persisted for up to 3 weeks after relief.

Chronic I/R injury induced NVCs in the cystometrogram in our study. Oxidative stress results in the generation of free radicals and oxidative damage [17]. Chronic oxidative stress may damage intrinsic nerves, resulting in partial denervation of smooth muscle. Denervation of the detrusor muscle leads to sensitization of the afferent pathway through postjunctional supersensitivity with increased neurotransmitters and upregulation of neurokinin receptors [18]. Denervation and nerve degeneration in sensory pathways during chronic I/R injury also cause an increase in the nerve growth factor level that may induce bladder hypersensitivity [19]. Additionally, chronic oxidative stress leads to the accumulation of calcium in the intracellular medium and the formation of metabolic end products that damage the detrusor musculature [17]. The resulting neurogenic and myogenic damage may cause detrusor overactivity but impaired contractility, resulting in NVCs in the cystometrogram.

Systemic administration of antioxidants reduced I/R injury of the bladder after relief. Several studies have proven that antioxidants reduce oxidative stress. Antioxidants have shown preventive and therapeutic benefits in several cardiovascular diseases [11]. Free radical scavengers decrease blood pressure and improve vascular function via biochemical mechanisms, such as normalizing the increased renal sympathetic nerve activity, plasma norepinephrine levels, and angiotensin type I receptor expression and enhancing carotid body chemoreceptor sensitivity to hypoxia [20]. The use of antioxidants after liver transplantation attenuates the effects of I/R-related oxidative stress and reduces lipid peroxidation [10]. However, few reports have shown that antioxidants can reduce oxidative stress in the bladder. I/R injury after BPH surgery has drawn increasing attention with the popularity of the HoLEP procedure, which can completely resect the adenoma. Determining whether the delivery of stable antioxidants to the target organ is possible is important at the time of systemic administration of antioxidants. We confirmed that delivery of effective antioxidants to the bladder was possible in our study. However, studies on bladder-specific delivery may be necessary in the near future.

Apoptosis due to I/R injury was mainly observed in the mucosal layer rather than in the muscle layer. Sensory neurons in the lamina propria, which are susceptible to I/R injury, may be irreversibly damaged by I/R injury [21]. Degeneration of sensory neurons in the mucosal layer may contribute to detrusor instability. In our study, more NVCs occurred in the untreated rats, which were not given antioxidants and sustained more I/R injuries. This finding could be inferred from the influence of apoptosis in the mucosal layer. This finding of our study suggested that the pathophysiology of an overactive bladder may be due to the neuroplasticity of peripheral sensory neurons rather than the myogenic hypothesis.

A difference in beta-3 adrenoceptor expression was observed depending on the degree of I/R injury. In our study, the administration of antioxidants for 3 week prevented the reduction of beta-3 adrenoceptor protein expression compared to the expression level in the untreated group. There are recent studies that the expression of beta-3 adrenoceptor mRNA may be dependent on the degree of obstruction. However, the results are still controversial. In the severe BOO group, the expression level of beta-3 adrenoceptor mRNA in the mucosa of prostatic urethra was significantly lower than that in the mild BOO group [22]. In contrast, pBOO has been reported to increase beta-3 adrenoceptor mRNA expression in bladder of rat models [23]. Activation of beta-3 adrenoceptor in the urothelium induces relaxation of the bladder detrusor muscle [24]. Clinically, some patients exhibit no effect of beta-3 agonists. The difference in the effect of beta-3 agonists may be explained by the differences in beta-3 adrenoceptor expression in the urothelium of the bladder.

## Conclusions

I/R injury that occurs after relief of pBOO causes histological and functional changes in the bladder. Apoptosis in the urothelium and the deposition of collagen fibers in the detrusor muscle layer were significantly increased. The incidence of NVCs was also increased. Additionally, the beta-3 adrenoceptor was associated with bladder overactivity caused by I/R injury. In our study, free radical scavengers prevented this oxidative stress. These preventive effects persisted for up to 3 weeks after relief.

